# On anammox activity at low temperature: effect of ladderane composition, process conditions and dominant anammox population

**DOI:** 10.1101/2019.12.15.873869

**Authors:** V Kouba, K Hurkova, K Navratilova, D Vejmelkova, A Benakova, M Laureni, P Vodickova, T Podzimek, P Lipovova, L van Niftrik, J Hajslova, MCM van Loosdrecht, DG Weissbrodt, J. Bartacek

## Abstract

The application of partial nitritation-anammox (PN/A) under mainstream conditions can enable substantial cost savings at wastewater treatment plants (WWTPs), but how process conditions and cell physiology affect anammox performance at psychrophilic temperatures below 15 °C remains poorly understood. We tested 14 anammox communities, including 8 from globally-installed PN/A processes, for (i) specific activity at 10-30 °C (batch assays), (ii) composition of membrane lipids (U-HPLC-HRMS/MS), and (iii) microbial community structure (16S rRNA gene amplicon sequencing). Crucially, the key parameters impacting anammox activity were the membrane lipid composition and cultivation temperature. The size of ladderane lipids and the content of bacteriohopanoids were key physiological drivers of anammox performance at low temperatures. Higher contents of (i) short C18 [3]-ladderane alkyl and (ii) large phosphatidylcholine headgroup were determined in anammox more active at 15-30 °C and 10-15 °C, respectively. At below 15 °C, the activation energies of most mesophilic cultures severely increased while those of the psychrophilic cultures remained stable; this indicates that the adaptation of mesophilic cultures to psychrophilic regime necessitates months, but in some cases can take up to 5 years. Interestingly, biomass enriched in the marine genus “*Candidatus* Scalindua” displayed exceptionally highest activity at 10-20 °C (0.50 kg-N.kg-VSS^−1^.d^−1^ at 10 °C, Ea10-30 °C = 51±16 kJ.mol^−1^), indicating outstanding potential for nitrogen removal from cold streams. Collectively, our comprehensive study provides essential knowledge of cold adaptation mechanism, will enable more accurate modelling and suggests highly promising target anammox genera for inoculation and set-up of anammox reactors, in particular for mainstream WWTPs.

**Highlights:**

- Ladderane size and cold exposure affected anammox activation energy (Ea).
- Ea improved with more C18 [3]-ladderanes over C20 and larger polar headgroup.
- Long-term cold exposure reduced Ea at 10-15 °C, not activity *per se*.
- Marine “*Ca*. Scalindua” was exceptionally suitable for cold streams.
- Anammox Ea at 15-30 °C was 79±18 kJ.mol^−1^.

**Graphical abstract:** 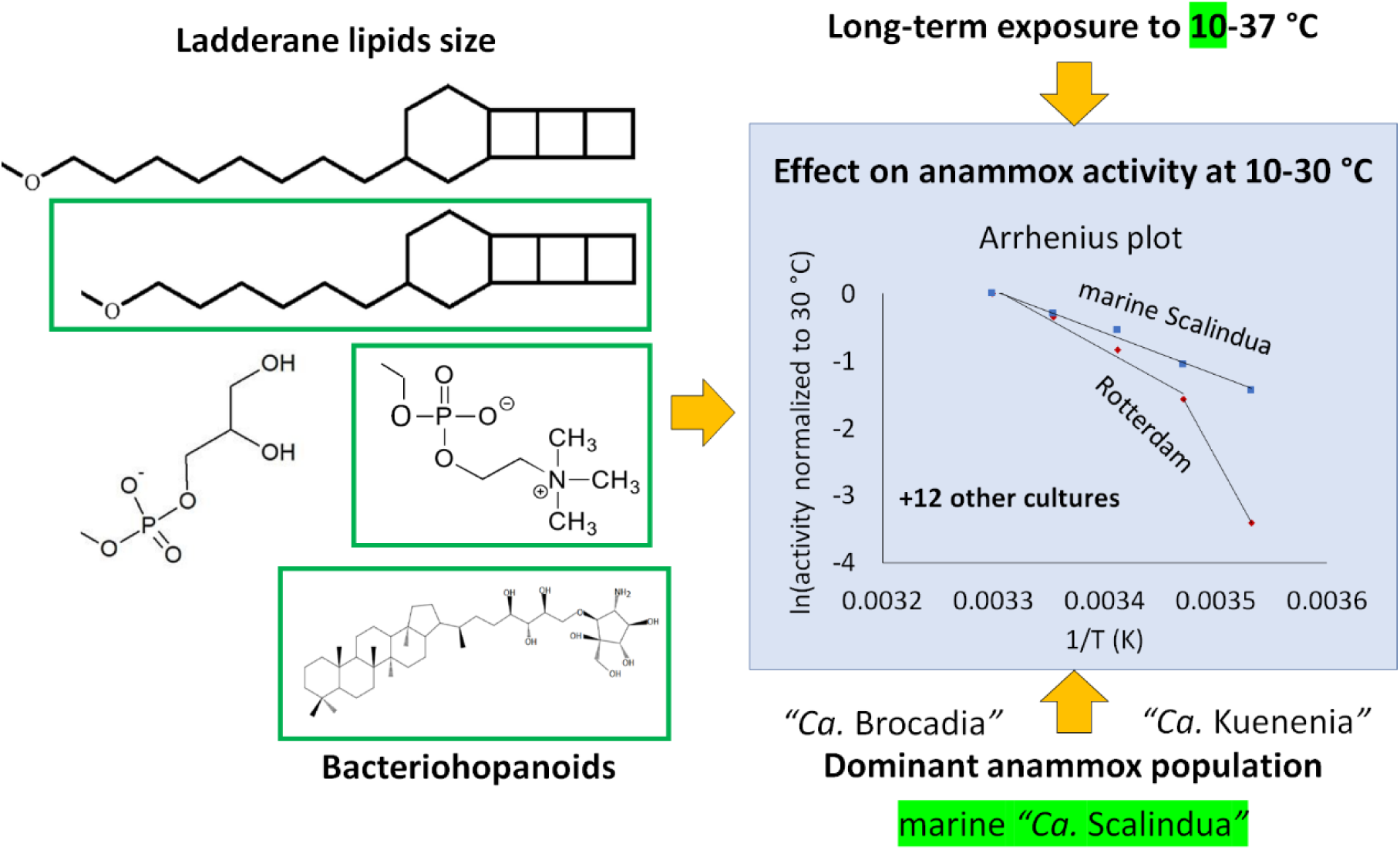

## 1 Introduction

Anaerobic ammonium oxidation (anammox) is an established microbial process for nitrogen removal from reject water (side streams) from sludge digestion and other nitrogen-rich and warm wastewaters. Compared to nitrification-denitrification, it does not require any exogenous organic carbon consumption and produces by up to 80% less excess sludge due to the autotrophic nature of anammox microorganisms. Because just 57% of the ammonium is oxidized to nitrite only, the combination of anammox with partial nitritation saves more than 50% in aeration energy and aeration system capacity (Daigger 2014, Jetten et al. 1997, Wett et al. 2007). According to Lackner et al. (2014), Bowden et al. (2015) and our own research, this technology has been implemented at over 150 full-scale anammox installations world-wide for the treatment of concentrated side streams, making side-stream anammox an established technology. At these installations, the parameters beneficial for the anammox process are high temperatures 30-37 °C and high concentrations of hundreds of mg of ammonium nitrogen per liter. A decrease in these process parameters unfavorably impacts process efficiency (Lackner et al. 2014). At 30-37 °C and an order of magnitude lower ammonium nitrogen, anammox has been reported to spontaneously occur in the more diluted main stream of municipal wastewater treatment plants (Cao et al. 2017a). This indicated that even a low ammonium concentration is not a bottleneck. In nature, anammox bacteria were detected in both marine and freshwater mesophilic (25-38 °C) and psychrophilic (10-25 °C) ecosystems (Wan et al. 2019, Wang et al. 2019, Zhao et al. 2019). This supports the potential to extend the applicability of anammox to the mainstream of wastewater treatment plants (WWTPs).

Currently, the main challenge in anammox research is its implementation in colder main-stream conditions, one of the main bottlenecks being the low activity of anammox bacteria at low temperatures (Cao et al. 2017b, Hoekstra et al. 2018, Seuntjens et al. 2016). This implementation will reduce operational and capital expenses (*i*.*e*., capacity of aeration system) for the removal of nitrogen from colder mainstream WWTPs and enable a more complete utilization of organic carbon in wastewater, *e*.*g*., for energy generation (Hejnic et al. 2016). Specifically, the low activity is cumbersome in psychrophilic main-stream reactors inoculated with mesophilic anammox cultures (Cao et al. 2017b). Per recent evidence, anammox can overcome cold stress and improve activity at low temperatures. This can result from gradual acclimation (De Cocker et al. 2018), enrichment of cold-adapted species (Hendrickx et al. 2014) or cold shocks (Kouba et al. 2017).

Nonetheless, the following questions still waits to be answered. The activity of anammox cultures as a function of temperature has yet to be reported in sufficient detail, such as for anammox genera other than “*Candidatus* Brocadia” and biomasses operated for the long-term in psychrophilic regime. In the few studies available, the effect of temperature on the activity of anammox cultures has been assessed using only a single genus (“*Ca*. Brocadia”), and biomass from either few mesophilic side-stream reactors (Lotti et al. 2014) or marine environments (Zhou et al. 2017). A proteomic study by Lin et al. (2018) has suggested that low temperatures do affect “*Ca*. Kuenenia” and “*Ca*. Jettenia” more strongly than “*Ca*. Brocadia”. Some other recent studies have associated anammox cold adaptation with an increased anammox activity and a shift in dominant anammox populations (Akaboci et al. 2018, De Cocker et al. 2018, He et al. 2018, Li et al. 2018, Wang et al. 2018b, Zhang et al. 2018a, Zhang et al. 2018b). Therefore, the currently rare cultures of anammox bacteria operated under a long-term psychrophilic regime may be affected by temperature differently than mesophilic cultures, since they will be dominated by cold-adapted microbial species, or by different species altogether. However, as the psychrophilic cultures have been made available only recently, they have never been characterized to sufficient detail. Specifically, correlations between anammox activities (in terms of absolute activities and activation energies) and long-term cultivation temperature and microbial community structure are yet to be addressed in a comprehensive survey.

One of the hypothetical mechanisms responsible for anammox adaptation to cold stress is the altered composition of ladderane phospholipids. Ladderanes are unique to anammox, likely reducing the diffusion of protons from anammoxosome organelle, thus enabling the slow anammox reaction (Moss et al. 2018). They consist of three or five concatenated cyclobutene rings bound to a polar head group by an ester or ether bond (Sinninghe Damsté et al. 2002). Generally, cold-adapted bacteria tend to synthesize more branched, unsaturated and shorter fatty acid phospholipids, so that their cytoplasmic membrane remains fluid, thus maintaining function of membrane proteins (Beales 2004). Only one single study on ladderanes in cold anammox bacteria has been published, suggesting that anammox cultivated at lower temperatures had contained more C18 than C20 ladderanes (Rattray et al. (2010). Their study also reported the ladderane composition of anammox cultures from multiple environments and WWTPs. However, correlating ladderane composition to culture activity as a function of temperature has yet to be investigated.

This study assessed the effect of temperature (10, 15, 20, 25, 30 °C) on the activity of 14 anammox biomasses originating from a representative set of full-scale reactors, pilot- and lab-scale models and highly enriched lab-scale cultures. The activities and activation energies were correlated with ladderane content, dominant anammox populations, cultivation temperature regime and relevant process conditions (*i*.*e*., one- or two-stage PN/A, cultivation of anammox in granules/flocs/carriers/free cells). Collectively, our findings identified the most suitable inocula and process conditions applicable for side- and mainstream anammox. They provide essential insights for process acclimation under various temperature regimes. While suggesting mechanisms anammox bacteria use to adapt to low temperature, the activation energies measured can sustain integration into mathematical models for process anticipation.

## 2 Materials and methods

### 2.1 Anammox biomasses

The mesophilic biomasses sampled from full-scale installations (Landshut / DE, Plettenberg / DE, Malmö / SE, Strass / AT, Tilburg / NL, Rotterdam / NL), enrichment of Kuenenia (Delft / NL) and Brocadia (Nijmegen / NL) and psychrophilic cultures from full scale WWTPs (Eisenhüttenstadt / DE, Xi’an / CN), pilot and laboratory reactors (Lemay / FR, Dübendorf1 / CH, Dübendorf2 / CH) and enrichment Scalindua (Nijmegen / NL) are described in Table 1.

**Table 1:** Description of tested anammox cultures named based on their original location, except for enrichments.

| ID | Description of technology | Influent | Sludge character | t (°C) | N-load (kg.m<sup>-3</sup>.d<sup>-1</sup>) | aeration / DO (mg.L<sup>-1</sup>) |
| --- | --- | --- | --- | --- | --- | --- |
| Full-scale installations |  |  |  |  |  |  |
| Eisenhüttenstadt / DE | SBR DEMON®, one-stage | reject water | flocs/granules | 21 | - | intermittent/- |
| Landshut / DE | TERRAMOX®, two-stage | reject water | flocs | 32 | 0.5 | no |
| Malmö / SE | AnitaMOX™, Anox Kaldnes, two-stage | reject water | biofilm, carriers K5** | 30 | 1.0 | no |
| Plettenberg / DE | SBR DEMON®, one-stage | reject water | flocs/granules | 30 | 0.3 | intermittent/- |
| Strass / AT | continuous DEMON®, one-stage | reject water | flocs/granules | 30 | 0.8 | intermittent/0.4 |
| Rotterdam / NL | ANAMMOX®, two-stage | reject water | granules | 37 | 3.4 | no |
| Tilburg / NL | ANAMMOX®, one-stage | reject water, CAMBI | granules | 37 | 1.3 | continuous/3.0-3.5 |
| Xian / CN | full-scale WWTP, activated sludge process | municipal sewage | biofilm, carriers | n.a. | n.a. | n.a. |
| Pilot and lab-scale systems |  |  |  |  |  |  |
| Lemay / FR | lab-scale 1.6 m<sup>3</sup>, VERI*, one-stage | pre-treated municipal sewage | biofilm, carriers K5** | 20 | 0.2-0.3 | continuous/0.7 |
| Dübendorf1 / CH | pilot-scale MBBR, EAWAG | municipal sewage pre-treated in A-stage | biofilm, carriers | 14 | n.a. | n.a. |
| Dübendorf2 / CH | lab-scale MBBR, EAWAG | synthetic | biofilm, carriers | 10 | n.a. | n.a. |
| Enrichments |  |  |  |  |  |  |
| Brocadia | enrichment, laboratory of Microbiology, Radboud University, NL | synthetic | granules | 30 | 0.36 | no |
| Kuenenia | enrichment, laboratory of EBT, TU Delft, NL | synthetic | planktonic | 30 | 1 | no |
| Scalindua | enrichment, laboratory of Microbiology, Radboud University, NL | synthetic | flocs | 20 | 0.93 | no |

### 2.2 Experimental set-up

The batch experiments were initiated by transferring anammox biomass to two reactors with a working volume of 1 L. The anammox biomass amount was set so that the duration of test at 30 °C was at least 45 min to allow for collection of a minimum of 4-5 samples, and the resulting biomass content was kept consistent during all temperature tests. The biomass in vessels was gently mixed by magnetic stirrers Heidolph MR Hei-Mix L at 250 rpm. To maintain vessel temperature at 5, 10, 15, 20, 25 or 30 °C, the vessels were cooled or heated using thermostats Julabo F12 (Julabo GmbH, Germany). Anoxic conditions were maintained by a gentle flushing of the headspace with dinitrogen gas (grade 4.0) at 50-200 mL.min^−1^. After the suitable temperature was established, pH was adjusted to 7.40±0.05 by 0.05 mol.L^−1^ HCl and NaOH. Then, NaNO_2_ and NH_4_Cl dissolved in 5-10 mL of tap water was dosed to both reactors, so that each assay started at 40 mg-N.L^−1^ of nitrite and at least 40 mg-N.L^−1^ of ammonium. The tests were done in duplicates, achieving a relative average deviation of 9.3±8.2%.

Regular sampling of batch reactors was carried out to analyse total ammonium nitrogen (TAN) as the sum of N-NH_3_ and N-NH_4_^+^, N-NO_3_^−^, N-NO_2_^−^, and to measure suspended and total solids concentration according to APHA (2005). The anammox activity was determined as a sum of nitrite and ammonium nitrogen removal rate, each calculated as a linear slope of nitrogen concentration development during respective batch tests. To avoid error due to changing affinity to substrate, only concentrations in the linear range were included.

The contribution of denitrification to nitrogen removal was calculated as the removal rate of nitrite higher than according to the anammox stoichiometry given by Strous et al. (1998).

To evaluate the effect of temperature on anammox activity, the activities were normalized to 30 °C. The activation energies were calculated according to the Arrhenius’ empirical model and its linearized version (equations 1 and 2), where *k* is the ratio of anammox activities at the lower (numerator) and higher (denominator) compared temperatures, *ln* is the natural logarithm, *A* is a constant pre-exponential factor, *Ea* is the activation energy (J.mol^−1^), *R* is the ideal gas constant (J.mol.^−1^.K^−1^) and T is the thermodynamic temperature (K). The data were linearized using equation 2, yielding either one or two *Ea*. Two *Ea* were chosen when the resulting correlation coefficient R^2^ was higher by 0.4. The exception was biomass Tilburg, where individual activation energy was attributed to each temperature interval of 10-15, 15-20, 20-25, and 25-30 °C.

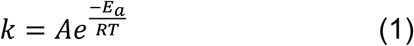

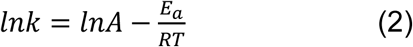

### 2.3 Analysis of bacterial community compositions by amplicon sequencing

All biomasses were analysed for their bacterial community compositions by 16S rRNA gene amplicon sequencing of distinct regions (16S V4 / 16S V3 / 16S V3-V4 / 16S V4-V5, 18S V4 / 18S V9, ITS1 / ITS2, Arc V4). Genomic DNA was extracted using the DNeasy® PowerBiofilm® Kit (MO BIO GmBH, Germany) following the manufacturer’s protocol and submitted to Novogene (Hong Kong, PRC) for amplicon sequencing using the MiSeq workflow (Illumina, US). Details on the method are described in the section Amplicon Sequencing Methodology of the Supporting Information.

### 2.4 Ladderane analysis

We used U-HPLC–HRMS/MS which is exceptionally sensitive and provided insight into the number of carbon atoms of these lipids. Per Rattray et al. (2008), the polar headgroup ionization differs substantially, so the results can be characterized only qualitatively.

#### 2.4.1 Reagents and chemicals

Deionized water was obtained from a Milli-Q® Integral system supplied by Merck (Darmstadt, Germany). HPLC-grade methanol, isopropyl alcohol, formic acid and ammonium formate (purity ≥ 99 %) were purchased from Sigma-Aldrich (St. Luis, MO, USA).

#### 2.4.2 Sample preparation

To extract ladderane phospholipids, a mixture of MeOH:DCM:10 mM ammonium acetate (2:1:0.8, v/v/v) was chosen according to Lanekoff & Karlsson (2010). Lyophilized anammox cultures were weighted (0.2 g) into a plastic cuvette and automatically shaken for 2 min with 2 mL of extraction solvent. The suspensions were sonicated for 10 min, centrifuged (5 min, 10000 rpm, 5 °C). Finally, 1 mL of supernatant was transferred into the vial before further analysis by ultra-high performance liquid chromatography coupled to high-resolution tandem mass spectrometry (U-HPLC– HRMS/MS).

#### 2.4.3 Ultra-high performance liquid chromatography coupled to high-resolution mass spectrometry (U-HPLC-HRMS)

The Dionex UltiMate 3000 RS U-HPLC system (Thermo Fisher Scientific, Waltham, USA) coupled to quadrupole-time-of-flight SCIEX TripleTOF^®^ 6600 mass spectrometer (SCIEX, Concord, ON, Canada) was used to analyse ladderane phospholipids. Chromatographic separation of extracts was carried out using U-HPLC system, which was equipped with Acquity UPLC BEH C18 column, 100Å, 100 mm × 2.1 mm; 1.7 µm particles (Waters, Milford, MA, USA). The mobile phase consisted of (A) 5 mM ammonium formate in Milli-Q water:methanol with 0.1% formic acid (95:5 v/v) and (B) 5 mM ammonium formate in isopropyl alcohol:methanol: Milli-Q water with 0.1% formic acid (65:30:5, v/v/v).

The following elution gradient was used in positive ionization mode: 0.0 min (90% A; 0.40 mL min-1), 2.0 min (50% A; 0.40 mL min^−1^), 7.0 min (20% A; 0.40 mL min^−1^), 13.0 min (0% A; 0.40 mL min^−1^), 20.0 min (0% A; 0.40 mL min^−1^), 20.1 min (95% A; 0.40 mL min^−1^), 22.0 min (90% A; 0.40 mL min^−1^).

The sample injection volume was set at 2 μL, the column temperature was kept constant at 60 °C and autosampler temperature was permanently set at 5 °C. A quadrupole-time-of-flight TripleTOF® 6600 mass spectrometer (SCIEX, Concord, ON, Canada) was used. The ion source Duo Spray™ with separated ESI ion source and atmospheric-pressure chemical ionization (APCI) was employed. In the positive ESI mode, the source parameters were set to: nebulizing gas pressure: 55 psi; drying gas pressure: 55 psi; curtain gas 35 psi, capillary voltage: +4500 V, temperature: 500 °C and declustering potential: 80 V.

The other aspects of the methodology were consistent with Hurkova et al. (2019), except of confirmation of compound identification, which used accurate mass, isotopic pattern and MS/MS characteristic fragments.

## 3 Results

### 3.1.1 All biomasses were dominated by “*Ca*. Brocadia”, except enrichments

#### 3.1.2 Bacterial community compositions by 16S rRNA gene amplicon sequencing

The main anammox genera detected across the lab-scale, pilot, and full-scale biomasses by amplicon sequencing were “*Ca*. Brocadia” (1 – 50%), “*Ca*. Scalindua” (0 – 11%) and “*Ca*. Kuenenia” (0 – 76%) (Fig. 1). Small relative abundances of “*Ca*. Anammoxomicrobium” were detected in several samples, less than 0.08% of relative abundance normalized to all bacteria OTU. Aside from the two enrichments (“*Ca*. Scalindua”, “*Ca*. Kuenenia”), “*Ca*. Brocadia” was the dominant anammox genus in all biomasses. Total anammox sequencing read counts relative to total bacteria varied from 1 – 78%. Detailed results of amplicon sequencing are described in supporting materials (Table S 1).

**Fig. 1:**
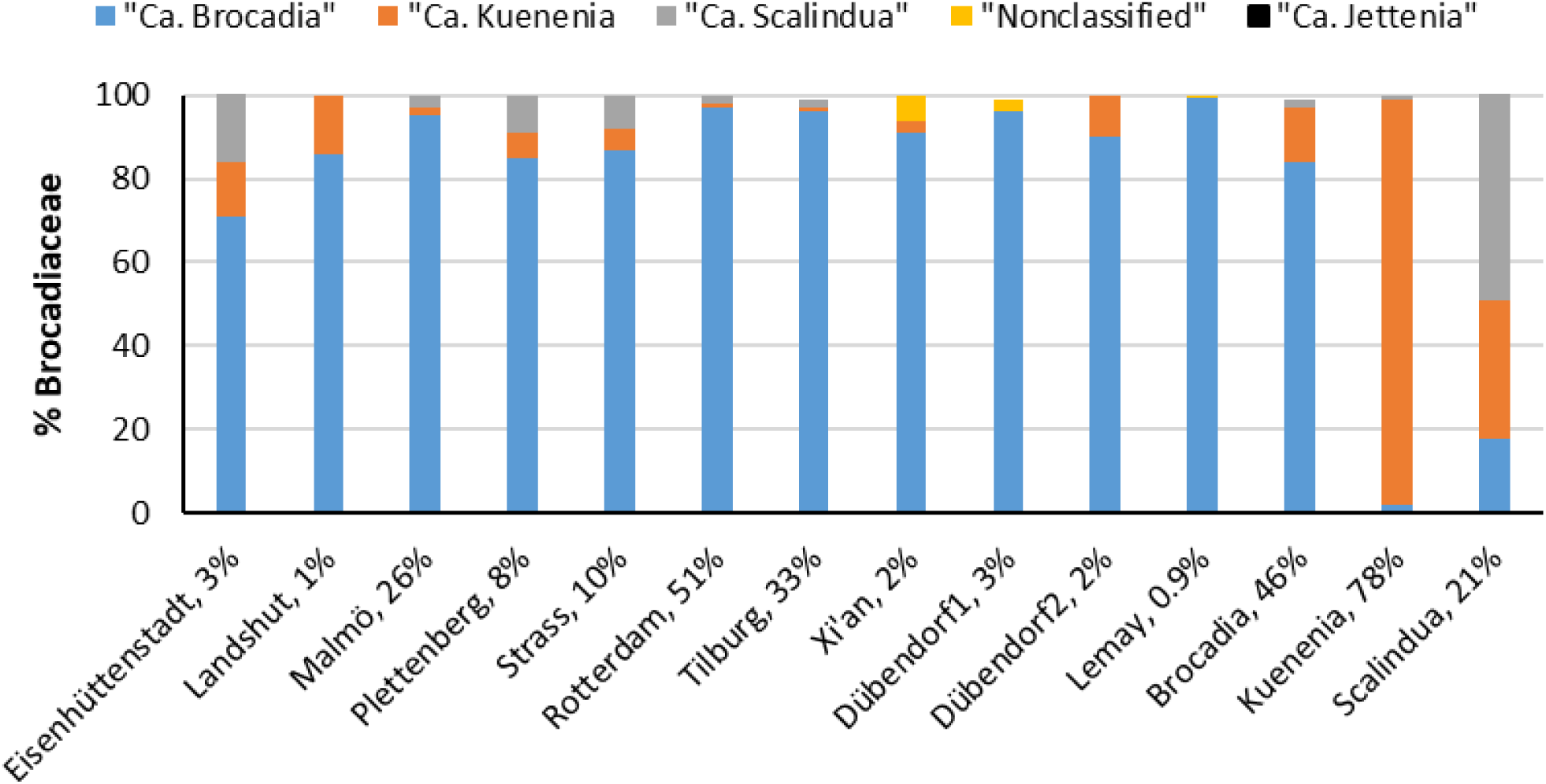
Relative abundances of all operational taxonomic units (OTUs) detected for each anammox genus within the anammox bacterial family of *Brocadiaceae*., The percentage of *Brocadiaceae* within the kingdom of Bacteria, as estimated by 16S rRNA gene-based amplicon sequencing analysis, is shown next to the name of the source of anammox culture. Aside from two enrichments, all biomasses were dominated by “*Ca*. Brocadia”.

#### 3.1.3 qPCR

qPCR provided a proxy for anammox abundance in the biomass, expressed as the ratio of anammox and the bacterial 16S rRNA genes. The efficiencies of the qPCR reaction for anammox and the bacterial 16S assays varied between 0.853-1.08 and 0.792-1.18, respectively. The ratios between the gene abundances varied from 0.02 to 0.41 (Fig. 2).

**Fig. 2:**
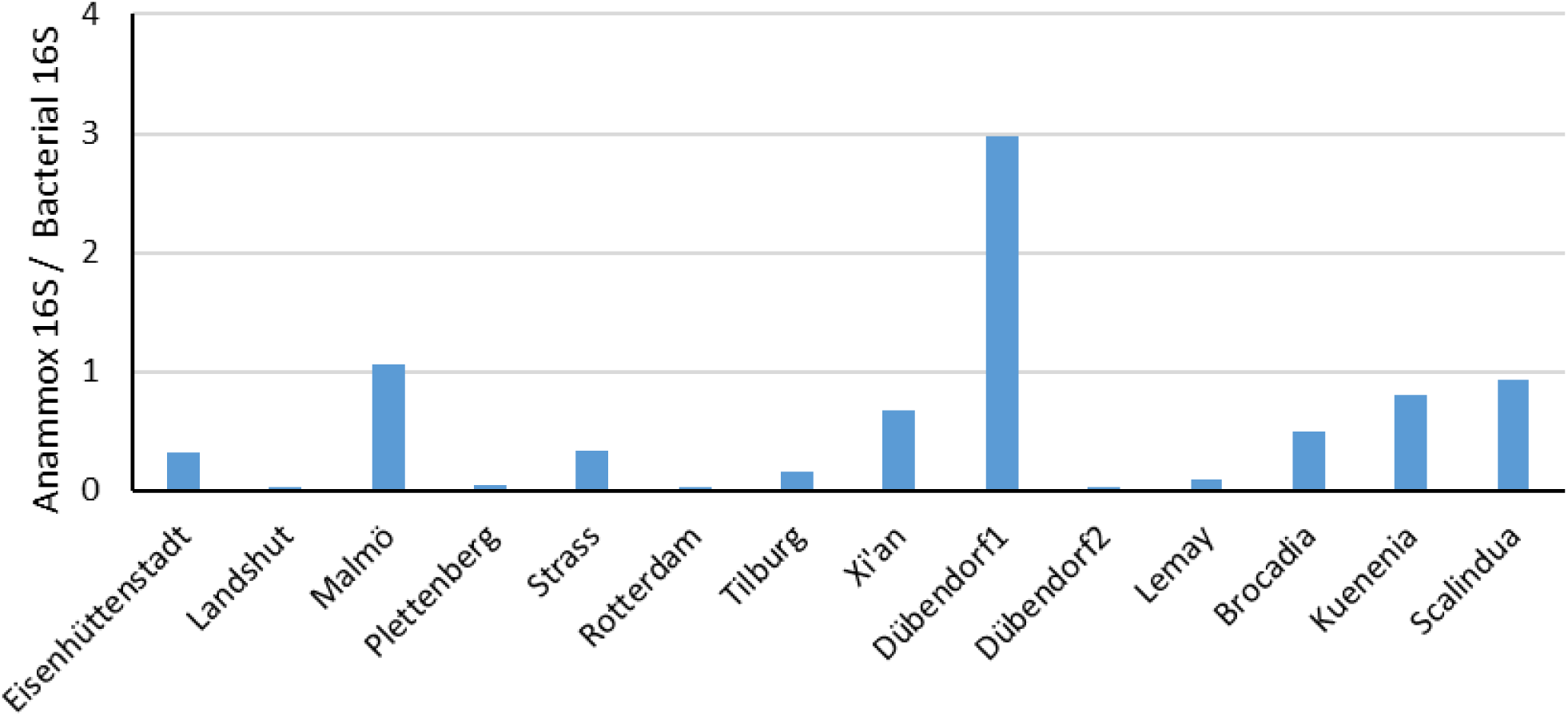
The ratio of abundance change of anammox-specific 16S rRNA genes and the total amount of bacterial 16S rRNA genes in anammox biomasses.

### 3.2 Effect of temperature on anammox performance: activity and activation energy (Ea)

The activity of various anammox cultures was expressed as the mass of ammonium and nitrite nitrogen metabolized per biomass weight (as volatile suspended solids, VSS) and time at 10-30 °C. As shown in Fig. 3, in the whole temperature range, the most active biomass of all was the marine enrichment of “*Ca*. Scalindua”. At 25-30 °C, similar activity was achieved by the enrichment of “*Ca*. Kuenenia”. Further, at 30 °C, the most active biomasses were Rotterdam (ANAMMOX®), Strass (DEMON®) and Malmö (AnitaMOX®). Following “*Ca*. Scalindua” and “*Ca*. Kuenenia” enrichments, these three mesophilic cultures (Rotterdam, Strass, Malmö) were also most active at 10 °C. Among psychrophilic cultures, the most active at 10 °C were the enrichment of “*Ca*. Scalindua”, followed by Lemay (0.031 kg-N.kg-VSS^−1^.d^−1^), Dübendorf1 (0.024 kg-N.kg-VSS^−1^.d^−1^) and Dübendorf2 (0.019 kg-N.kg-VSS^−1^.d^−1^).

**Fig. 3:**
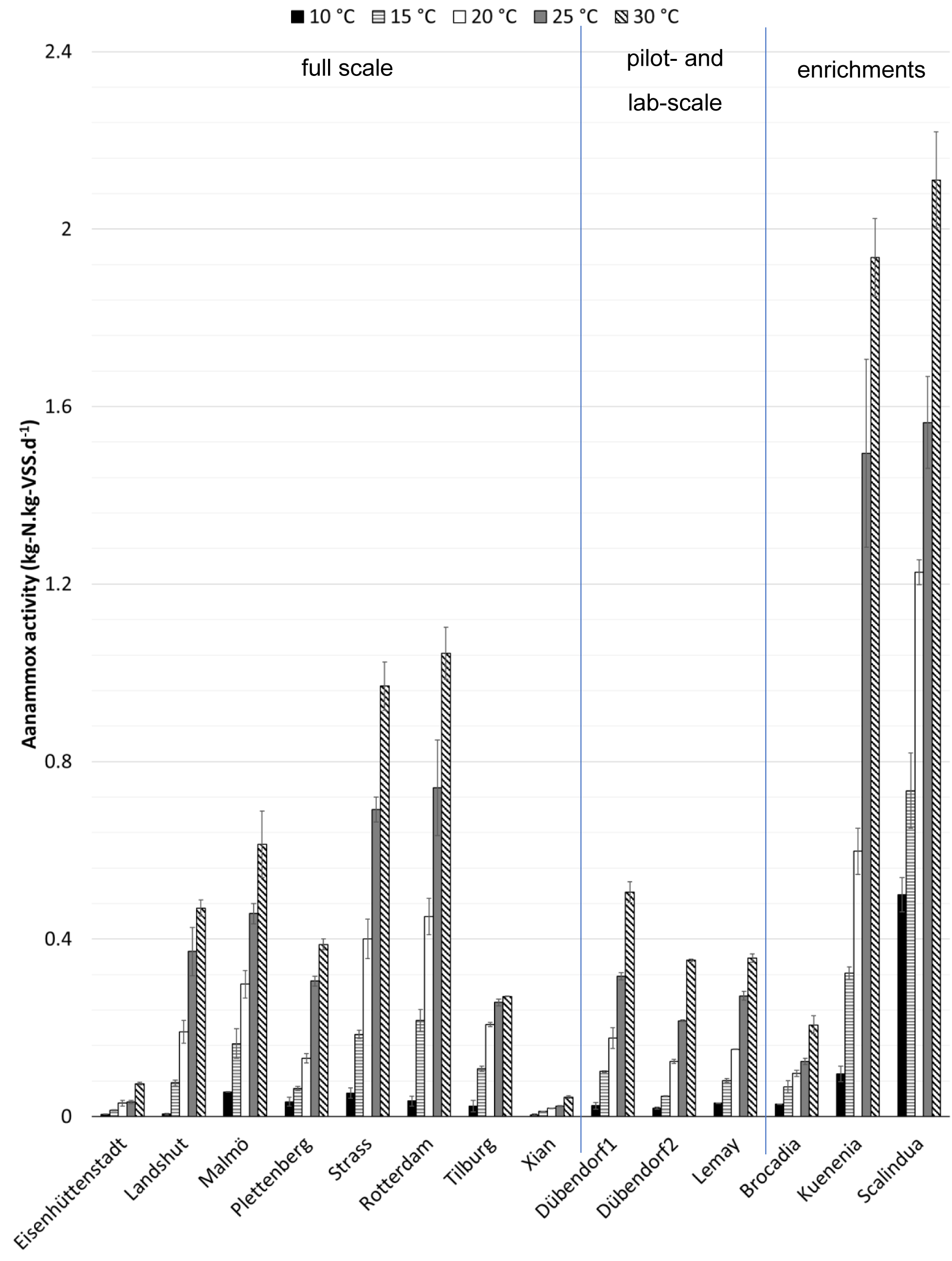
Effect of temperature on the specific anammox activity of multiple anammox biomasses from pilot and full scale installations and from laboratory enrichments (“*Ca*. Brocadia”, “*Ca*. Kuenenia”, “*Ca*. Scalindua”). “*Ca*. Scalindua” had the highest activity at 10-20 °C of all cultures.

To describe the effect of temperature on anammox biomasses, the activities were normalized at 30 °C and Ea was determined as a temperature coefficient for each culture (Table 2). At 15-30 °C, all anammox cultures were characterized by a similar Ea of 79±19 kJ.mol^−1^. All but one psychrophilic culture could be described by a single Ea over the range from 10-30 °C, similar to some mesophilic ones (enrichments “*Ca*. Kuenenia” and “*Ca*. Brocadia”, and biomass Plettenberg). Other mesophilic cultures were affected by temperature at 10-15 °C more severely (Table 2). Tilburg could only be described by a separate Ea for each of the four temperature intervals. Alternately, Tilburg and also some other cultures (*e*.*g*., Strass, Malmö) could be more accurately described by a quadratic function (Fig. S 1).

**Table 2:** Activation energies (Ea) of anammox cultures reported in the literature and determined in this study. The cultures are ranked from lowest to highest Ea under 10-15 °C, *i*.*e*., anammox cultures most adapted to low temperatures are on top.

| Reference | Character/culture origin, operational temperature/dominant anammox genera/species | t (°C) range for Ea | Ea (kJ.mol<sup>-1</sup>) |
| --- | --- | --- | --- |
| **This study** | **“Ca. Scalindua” enrichment, 20 °C** | **10-30** | **51±16** |
| (Rysgaard et al. 2004) | sediments, Greenland | -2-13 | 51 |
| (Dalsgaard and Thamdrup 2002) | sediments, Skagerrak, North Sea | 6.5-37 | 61 |
| Dosta et al. (2008) | biofilm, granules, “Ca. Kuenenia stuttgartiensis”, 30 °C, two-stage | 10-40 | 63 |
| (Puyol et al. 2014) | granules, “Ca. Brocadia fulgida”, 30 °C, two-stage | 10-30 | 64 |
| (Hoekstra et al. 2018) | free cells, MBR, 20-30 °C, “Ca. Brocadia fulgida”, two-stage | 15-30 | 64±28 |
| (Hendrickx et al. 2014) | flocculent sludge, “Ca. Brocadia fulgida”, 10 °C, two-stage | 5-17 | 66 |
| **This study** | **enrichment, “Ca. Brocadia anammoxidans”** | **10-30** | **71±36** |
| (Hendrickx et al. 2012) | granules, biofilm, “Ca. Brocadia fulgida”, 20 °C, two-stage | 10-20 | 72 |
| (Park et al. 2017) | biofilm, non-identified anammox or planktomycte bacteria, 20 °C, two-stage | 10-30 | 73 |
| **This study** | **Eisenhüttenstadt – suspension/granules, full scale, DEMON®, 21 °C, “Ca. Brocadia”** | **10-30** | **79±13** |
| **This study** | **Xi’an, main stream of WWTP, full scale, “Ca. Brocadia”** | **10-30** | **83±41** |
| **This study** | **enrichment, “Ca. Kuenenia stuttgartiensis”, 30 °C** | **10-30** | **83±42** |
| **This study** | **Lemay – moving bed biofilm reactor, K5 carriers, Veolia Research and Innovation, 1.6 m<sup>3</sup> reactor, 20 °C, “Ca. Brocadia”** | **10-30** | **86±3** |
| (Park et al. 2017) | granules, “Ca. Kuenenia stuttgartiensis”, 35 °C, two-stage | 10-35 | 89 |
| (Isaka et al. 2008) | encapsulated biomass, “Ca. Kuenenia stuttgartiensis”, “Ca. Brocadia anammoxidans”, Planctomycetes KSU-1 (AB057453), two-stage | 6-28<br>28-37 | 93<br>33 |
| **This study** | **Plettenberg – granules, full scale, DEMON®, 30 °C, one-stage, “Ca. Brocadia”** | **10-30** | **93±14** |
| **This study** | **Dübendorf2 – moving bed biofilm reactor (K5, Anox Kaldnes), 10 °C, “Ca. Brocadia”** | **10-30** | **104±32** |
| (Park et al. 2017) | biofilm, “Ca. Kuenenia stuttgartiensis”, “Ca. Brocadia caroliniensis”, “Ca. Brocadia fulgida” and other planktomycte bacteria, 20 °C, two-stage | 10-25 | 108 |
| (Lotti et al. 2014) | granules, 10 °C, “Ca. Brocadia sinica”, one-stage | 10-30 | 110 |

Table 2 (*continued*): Activation energies (Ea) of anammox cultures reported in the literature and determined in this study. The cultures are ranked from lowest to highest Ea under 10-15 °C, *i.e.*, anammox cultures most adapted to low temperatures are on top.
|  |  |  |  |
| --- | --- | --- | --- |
| (Lotti et al. 2014) | IC, full-scale, granules, "Ca. Brocadia fulgida", 30-35 °C | 10-30 | 117 |
| (Puyol et al. 2014) | flocs, "Ca. Brocadia caroliniensis", 30 °C, two-stage | 10-30 | 124 |
| (Lotti et al. 2014) | SBR (granules, 20 °C, "Ca. Brocadia fulgida"), one-stage | 10-15<br>15-30 | 140<br>52.3 |
| **This study** | **Malmö – moving bed biofilm reactor, K5 carriers, full scale, Anox Kaldnes, 30 °C, two-stage, "Ca. Brocadia"** | **10-15**<br>**15-30** | **150±10**<br>**63±12** |
| (Wang et al. 2018a) | granules, "Ca. Kuenenia Stutgartensis", two-stage | 10-20<br>20-33 | 153<br>9.4 |
| **This study** | **Strass – granules/flocs, full scale, DEMON®, 30 °C, one-stage, "Ca. Brocadia"** | **10-15**<br>**15-30** | **170±50**<br>**80±7** |
| **This study** | **Dübendorf1 – biofilm on MBBR, carriers Wabag Fluopur®, 14 °C, pilot-scale, one-stage, "Ca. Brocadia"** | **10-15**<br>**15-30** | **193**<br>**78±7** |
| (Lotti et al. 2014) | airlift, granules, "Ca. Brocadia sinica", 30-35 °C | 10-15<br>15-30 | 204<br>60 |
| **This study** | **Tilburg – granules, full scale, ANAMMOX®, 37 °C, one-stage "Ca. Brocadia"** | **10-15**<br>**15-20**<br>**20-25**<br>**25-30** | **207**<br>**92**<br>**31**<br>**8** |
| (Strous et al. 1999) | aggregate biomass, 32-33 °C, | 20-43 | 70 |
| **This study** | **Rotterdam – granules, full-scale, two-stage, ANAMMOX®, 37 °C, "Ca. Brocadia"** | **10-15**<br>**15-30** | **248**<br>**76±26** |
| (Lotti et al. 2014) | enrichment in MBR, free cells, "Ca. Brocadia fulgida", 30 °C | 10-15<br>15-30 | 325<br>94 |
| **This study** | **Landshut – flocs, full scale, TERRAMOX®, 32 °C, two-stage, "Ca. Brocadia"** | **10-15**<br>**15-30** | **360±140**<br>**89±20** |

\*unclear determination of Ea, arguable linear fit for biofilm (Fig 1 in Dosta et al. (2008); +calculated by the authors of this study based on data published in Lotti et al. (2014).

### 3.3 Ladderane composition

Membranes in original samples of anammox bacteria were investigated for the length of ladderane core lipids on *sn-1* and *sn-2* positions and polar head composition by U-HPLC-MS/MS. We detected one or two ether-bound C20-[5]-ladderanes; one of these positions were occupied by C20-[3]-, C18-[5]- and C18-[3]-ladderane ether or ester, in one case also a C22-[5]-ladderane ester, or a straight or branched alkyl chain with 14-16 carbon atoms. The polar head groups detected were either choline, ethanolamine or glycerol, mostly choline except for “*Ca*. Scalindua” enrichment (Fig. 4). Finally, the following triterpenoids were identified in anammox enrichments (ordered from lowest to highest abundance): squalene, bacteriohopanetetrol, and bacteriohopanetetrol cyclitol ether (for more details refer to Kouba et al., *submitted*).

**Fig. 4:**
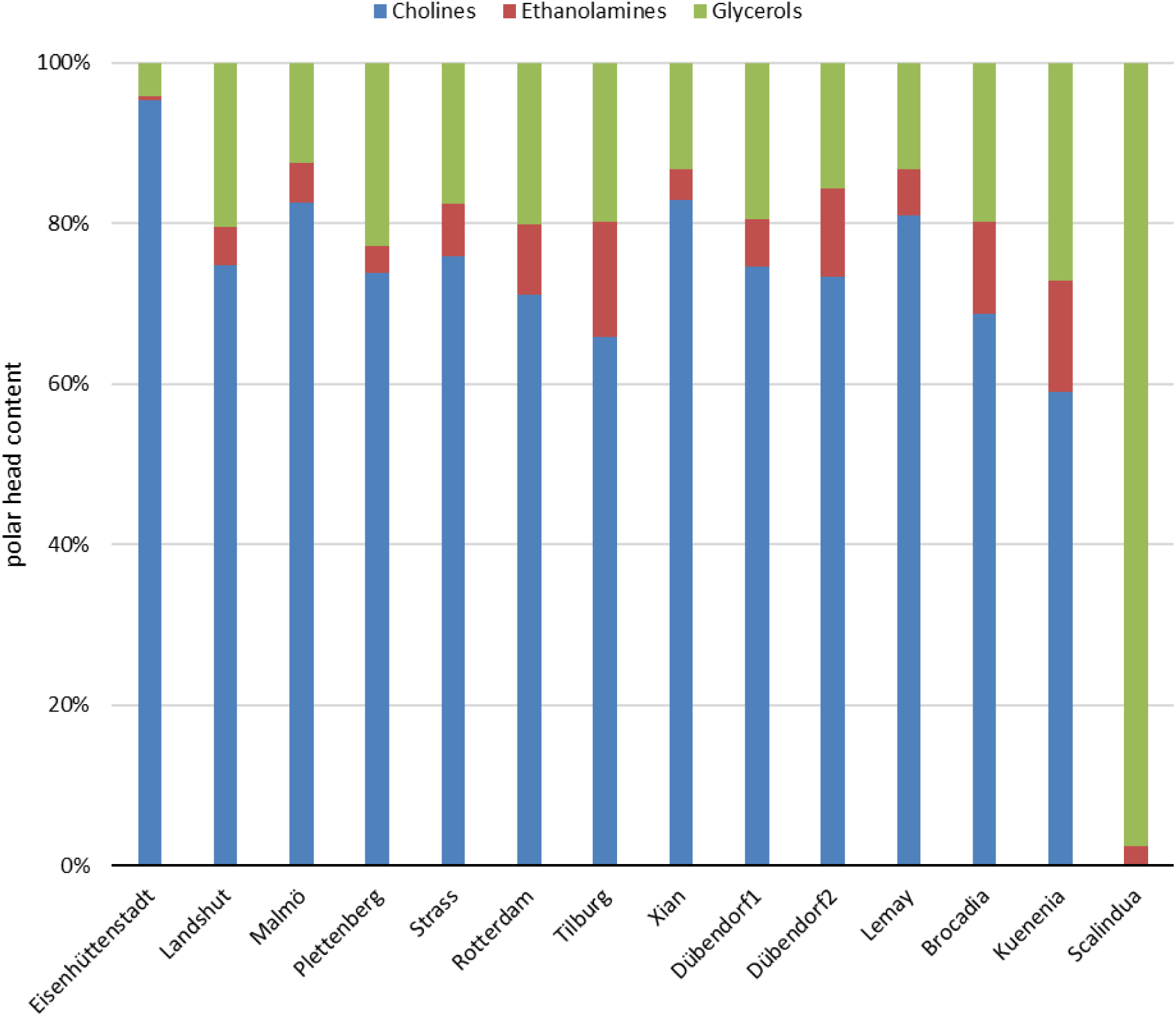
Polar head content of ladderane phospholipids in various anammox cultures. The ladderane phospholipid headgroup was mostly phosphatidylcholine in all cultures except of “*Ca*. Scalindua” enrichment.

## 4 Discussion

### 4.1 Cultivation temperature impacts Ea10-15

To date, literature provided only limited evidence on the long-term effect of low cultivation temperatures on the performance of originally mesophilic anammox cultures (Hoekstra et al. 2018, Wang et al. 2018b). Our results in combination with literature data show that a long-term cultivation at low temperature has a positive impact on the Ea of anammox conversion in the range of 10 to 15 °C (Fig. 5). Psychrophilic cultures had been operated under this regime for 8 months to 5 years. Across this timeframe, the operation length seemed neither to affect the anammox activation energy from 10 to 30 °C nor the activity from 10 to 20 °C. Thus, within these ranges, exposure to psychrophilic conditions for tens of months seems to be sufficient for inducing such adaptive response.

**Fig. 5:**
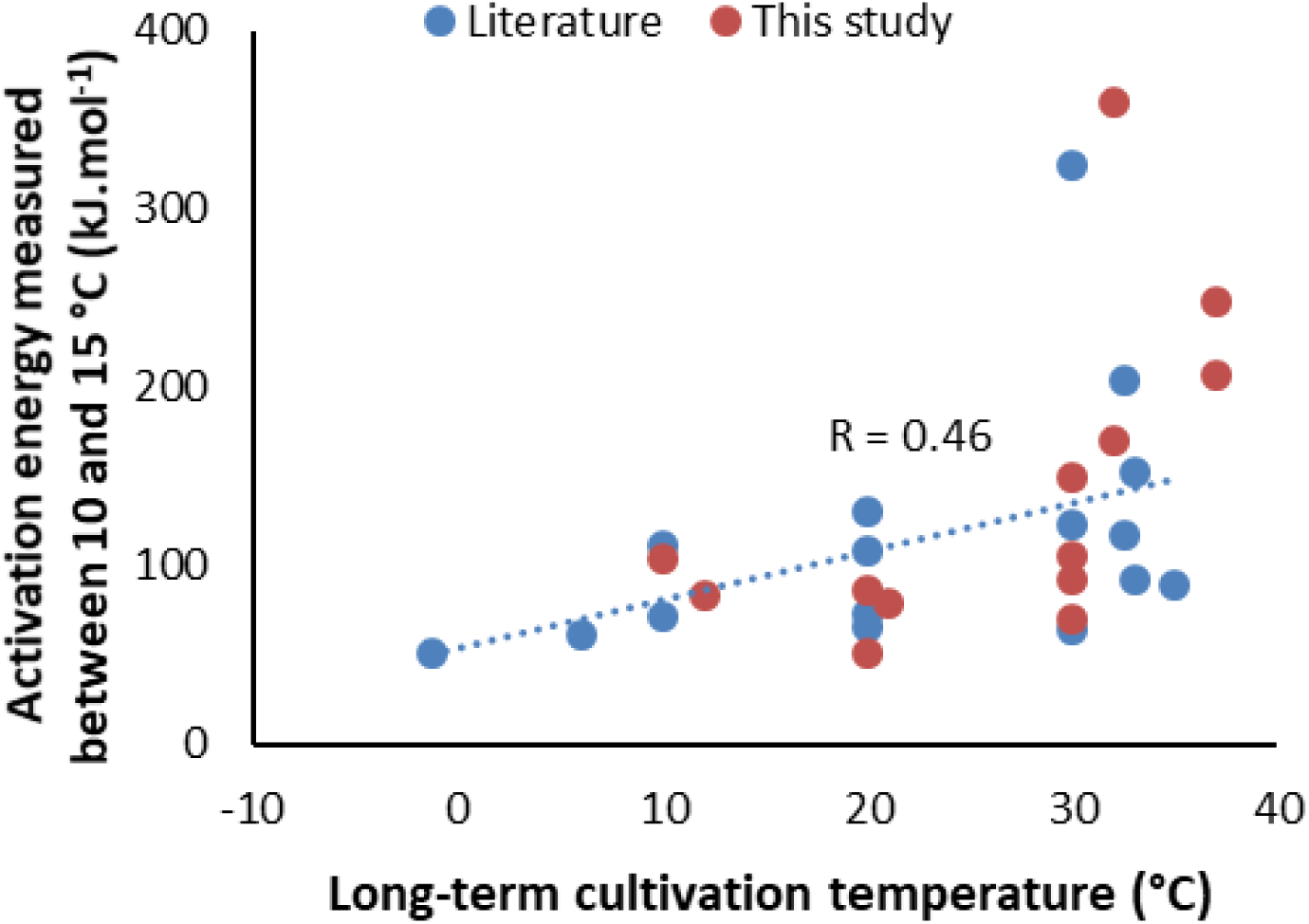
Correlation function of anammox long-term cultivation temperature and activation energy between 10-15 °C, including Pearson’s linear correlation coefficient (R=0.46), summarizing our and literature data. Activation energy at 10-15 °C of anammox biomasses cultivated in the mesophilic temperatures (30-37 °C) was in the range of 64-360 kJ.mol.^−1^, whereas the cultures operated at low temperatures up to 21 °C had substantially lower Ea (51-131 kJ.mol^−1^).

In terms of absolute activity at 10-15 °C, mixed cultures operated in the psychrophilic regime (not “*Ca*. Scalindua” enrichment) were not the most active (Fig. 3). Thus, our data do not confirm the hypothesis emitted by Lotti et al. (2014) that operation under psychrophilic temperatures improves the maximum anammox activities beyond values achieved by mesophilic cultures.

### 4.2 Implications for mathematical modelling of anammox activation energies

Most authors view 15 °C as a breaking point under which anammox bacteria may be more negatively affected by temperature (Cao et al. 2017b). This can be interpreted as if the effect of temperature on anammox activity as expressed by activation energy (Ea) is consistent throughout two ranges: 10-15 °C (hereafter referred to as Ea10-15) and 15-30 °C (Ea15-30). We show that this was true only for some mesophilic cultures. Therefore, to assume a single Ea10-15 for all mesophilic cultures is short sighted. For modelling purposes, Ea10-15 specific to the anammox biomass used should be obtained experimentally. However, almost all psychrophilic anammox cultures in this study could be described by a single activation energy for the whole range from 10 to 30 °C (Table 2), showing that there is no intrinsic difference in temperature response below 15 °C.

In our experiments at 15-30 °C, the anammox cultures had an activation energy of 79±18 kJ.mol^−1^ (average±standard deviation), which was identical to the average value 79±31 kJ.mol^−1^ calculated from the literature data (Table 2). In comparison, the standard activation energy for nitrification is similar 70 kJ.mol^−1^ (Wiesmann 1994). The activation energy range of mesophilic cultures at 10-15 °C was 71-360 kJ.mol^−1^.

### 4.3 Anammox performance of various genera: potential of marine “*Ca*. Scalindua”

The marine enrichment of “*Ca*. Scalindua” displayed the highest specific anammox activity at 10-20 °C. This was shown only in activities expressed per g-VSS (Fig. 3). In the literature, “*Ca*. Scalindua” is an organism that has mainly been recovered from marine environments (Cai et al. 2019, Zheng et al. 2019). We detected “*Ca*. Scalindua” also in biomasses treating supposedly less saline pre-treated sewage and reject water, but in relatively lower abundance compared to other genera, and those biomasses were not as active as the marine one. This study is the first to highlight such exceptional metabolic performance under 10-20 °C. Thus, we indicate that implementation of “*Ca*. Scalindua” to N-removal processes treating cold marine streams, and potentially cold streams in general, can be extremely beneficial. This should be considered when choosing appropriate inoculum and reactor design, however challenging.

The exceptional performance of “*Ca*. Scalindua” under low temperatures raises inquiry into their membrane physiology. As described in more detail in Kouba et al. (*submitted*), the [3]-ladderanes (Ea15-30 = 51 kJ.mol^−1^, C20/(C18+C20) [3]-ladderane ether = 0.11; Fig. 6) and [5]-ladderane esters alkyl moieties in “*Ca*. Scalindua” ladderane phospholipids were exceptionally short, having the highest relative content of C18 compared to C20 alkyls. Reduced length of phospholipid alkyls is typical in cold-adapted bacteria, as narrower membrane maintains its fluidity at lower temperature, thus maintaining function of membrane proteins (Siliakus et al. 2017). “*Ca*. Scalindua” also had an exceptionally high content of bacteriohopanoids, that were hypothesized to maintain membrane viscosity under cold stress (Boumann et al. 2009). In sum, the membrane of “*Ca*. Scalindua” appears exceptionally suitable to low temperatures.

**Fig. 6:**
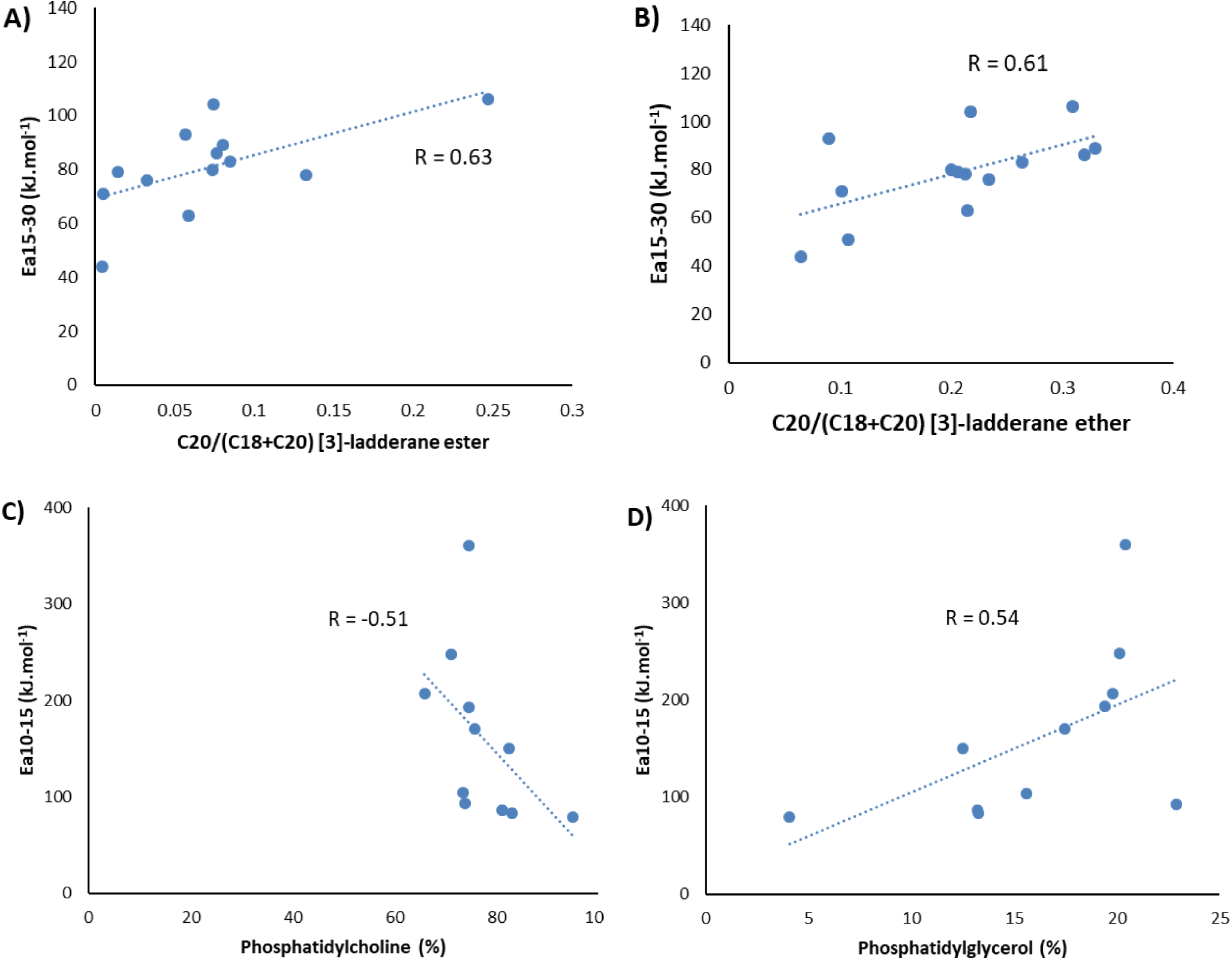
Correlation of anammox cultures ladderane phospholipid composition and activation energy between 15-30 °C and 10-15 °C, according to Pearson, summarizing our data. In A, *Ca*. Scalindua was excluded as outlier. Panel B contains all cultures. In C and D, only mixed cultures dominated by “*Ca*. Brocadia” are included.

Conversely, the polar headgroup of ladderane phospholipids was almost exclusively phosphatidylglycerol, while other cultures contained mostly phosphatidylcholine. Phosphatidylglycerol is smaller and thus less disruptive to the membrane packing which makes it less suitable for low temperatures (Siliakus et al. 2017).

Further, the adaptation to saline environment may induce the pre-disposition of “*Ca*. Scalindua” to low temperatures. Various species of “*Ca*. Scalindua” were reported to be adapted to low temperatures by ‘salt-in’ strategy, and early evidence points also to one species synthesizing compatible solutes (*i*.*e*., glutamate, glutamine, proline) (Speth 2016). Freezing point of aqueous solution decreases at elevated content of salts or these solutes, thus appearing as another mechanism making “*Ca*. Scalindua” especially cold-adapted.

### 4.4 Ladderane composition and anammox performance

Generally, in bacterial membranes, reducing temperature arranges lipids into more compact formations, thus reducing the membrane fluidity/flexibility. However, the membrane flexibility is crucial for the function of membrane proteins. Thus, bacteria maintain their membrane fluidity by synthesizing shorter, branched and unsaturated alkyl chains, larger polar headgroup and more terpenoids (Siliakus et al. 2017). However, the data on anammox membrane composition and anammox performance under low temperatures are missing, as the only closely related study restricted itself to the suggestion that cold anammox had more C18 compared to C20 [5]-ladderane esters, while the performance of such cultures remained untested (Rattray et al. 2010).

In our study, the crucial membrane features correlating to anammox activation energies at 10-30 °C were (i) the length of [3]-ladderanes and (ii) polar headgroup size. First, anammox cultures with higher content of C18 compared to C20 [3]-ladderanes chains had lower activation energy at 15-30 °C (Fig. 6). Interestingly, this did not appear to involve ladderanes with five concatenated cyclobutane rings, and not only ladderane esters as in Rattray et al. (2010), but also ethers. Second, in mixed cultures dominated by “*Ca*. Brocadia”, those with lower activation energy at 10-15 °C contained more phosphatidylcholine and less phosphatidylglycerol (Fig. 6). As choline is larger than glycerol or ethanolamine, it is thought to introduce additional disruption into membrane lipid packing (Siliakus et al. 2017). Similarly, a larger polar phospholipid head group has been shown to maintain membrane fluidity in barophilic bacteria (Jebbar et al. 2015). Overall, we provide the first evidence on the correlation between ladderane phospholipid composition and anammox activity.

Further, we detected higher content of monoalkyl ether phospholipids in cultures with higher activation energy at 10-15 °C (Fig. S 2). However, monoalkyl ether phospholipids and monoether could be not only one of the final membrane building blocks, but also perhaps an intermediate of lipid biosynthesis, or a by-product of cell lysis. Importantly, hydrocarbons and polar head attached to the glycerol backbone can be cleaved off by a phospholipase enzyme (Paltauf 1994).

Anammox cultures are known to contain triterpenoids such as various bacteriohopanoids (Rush et al. 2014) and squalene (Rattray et al. 2008), and these were suggested to play a role in maintaining membrane fluidity (Boumann et al. 2009). In contrast to ladderane lipids, these triterpenoids are not exclusively synthesized by anammox bacteria, so we analysed them only in highly enriched cultures, detecting bacteriohopanetetrol cyclitol ether, bacteriohopanetetrol and squalene in enrichments of “*Ca*. Scalindua”, “*Ca*. Brocadia”, and “*Ca*. Kuenenia”. Their signal intensity was correlated to activation energy at 10-30 °C, suggesting that their abundance may also contribute to maintaining anammox membrane fluidity at low temperatures.

The importance of shorter ladderane ethers, not only esters, suggests that future studies on anammox adaptation to different temperatures should not restrict their focus to any particular ladderane group. Rather, we advise a thorough assessment of the whole ladderane content, including ether-bound ladderane alkyl moieties.

### 4.5 Biomass growth mode and PN/A configuration

In all cultures, the biomass growth mode did not seem to correlate with anammox performance, with one exception. Free-cell anammox cultures consistently exhibited the highest maximum specific activity at 25-30 °C (Fig. S 2). This is probably due to the fact that planktonic cultures contain fewer non-anammox populations and less inactive organic matter (e.g. extracellular polymeric substances), both of which make them more active overall.

In the mixed psychrophilic cultures, the Lemay biomass (AnoxKaldnes K5 carriers) was six-fold more active than in the Xi’an (MBBR, main-stream) and Eisenhüttenstadt (granules/flocs, side-stream) cultures. We suspect that these less active cultures were exposed to more organic carbon in the municipal wastewater and flocculating agent, respectively. Thus, we hypothesize that their lower anammox activity may be due to stronger competition for nitrite from denitrifiers and that biomass growth mode may not have been the main factor. Nevertheless, certain biomass growth mode properties, such as biofilm depth, may impact substrate or toxin uptake rate, thereby affecting anammox performance.

### 4.6 PN/A configuration

Lotti et al. (2014) hypothesized that anammox growing in one-stage PN/A may become adapted to oxygen inhibition, and that the mechanism for maintaining homeostasis under oxygen inhibition may also alleviate cold stress. In this study, we did not detect a correlation between one or two-stage PN/A operation and anammox cultures performance at low temperature.

### 4.7 Anammox temperature optima

The temperature optimum of all the tested anammox cultures was ≥30 °C, including the long-term psychrophilic enrichments. In contrast, enrichments from arctic conditions had a much lower average optima of 12 °C (Rysgaard et al. 2004). Because some long-term lab-scale experiments at 10-20 °C observed temperature optima up to 25–30 °C, which is less than typically reported optima of 35-38 °C (Hendrickx et al. 2014, Hu et al. 2013, Park et al. 2017), we wondered whether longer exposure to a psychrophilic regime might reduce the optima even further. But this was not the case. Our psychrophilic cultures came from multi-year full-scale and lab-scale operations and contained a variety of anammox populations, including marine “*Ca*. Scalindua” enrichment, so their high temperature optima may be related to other factors, such as even lower cultivation temperatures (<10 °C, freeze-thaw cycles), or specific ladderane composition.

## 5 Conclusions

We have shown that, irrespective of the multiple conditions tested, the performance of anammox bacteria was crucially affected by both lipid composition and long-term exposition to low temperature. Most importantly, as anammox performance at low temperatures closely correlated with ladderane size and bacteriohopanoid abundance, these anammox membrane components proved to be key aspects of cold anammox physiology. Furthermore, long-term operation under psychrophilic conditions, while not always necessarily enhancing absolute activity, consistently improved the anammox temperature coefficient at 10–15 °C (85±49 kJ.mol^−1^, median±standard deviation). The activation energies of mesophilic cultures at 10–15 °C are highly diverse (160±95 kJ.mol^−1^), stressing the need for individual assessment of such cultures when modelling their activity. In addition, we showed the exceptional performance of a cold-adapted enrichment of marine “*Ca*. Scalindua”, highlighting its potential for nitrogen removal from cold and more saline streams, which is crucial when choosing the most appropriate inoculum and reactor set-up. Collectively, these findings, based on a complex assessment of metabolic activities, microbial community structure and membrane lipids in 14 anammox cultures and on a comprehensive literature survey, provide essential knowledge for the more accurate modelling for instance by the inclusion of measured activation energies, inoculation and set-up of anammox reactors, in particular for the main stream of WWTP.

## 6 Acknowledgement

The authors acknowledge the financial support of the Czech Ministry of Education Youth and Sports through project GACR 17–25781S, and internal partial funding from the TU Delft. Michele Laureni was supported by a Marie Skłodowska-Curie Individual Fellowship (MixAmox; 752992). The authors also thank the researchers, process engineers and operators for providing and/or maintaining the anammox biomass, namely Katinka van de Pas-Schoonen and Guylaine Nuijten (Microbiology, Radboud University), Albert Regiert (Stadtwerke Landshut, Germany), Martin Hell (AIZ, Strass im Zillertal, Austria), Jürgen Koepke (TAZV Oderaue, Germany), Jon Albizuri (Anox Kaldnes, Sweden), Romain Lemaire (Veolia Research and Innovation, France), Job Robben and Gijs Lavrijsen (Waterschap De Dommel, Netherlands), Jeroen van Waveren (Waterschap Hollandse Delta, Netherlands), Hans-Joachim Hölter (Ruhrverband, Germany), Xiaochang Wang and Jin Pengkang (Xi’an University of Architecture and Technology, China) and Adriano Joss and Damian Hausherr (Eawag, Switzerland). The authors also acknowledge the contribution of Craig Alfred Riddell in editing the manuscript. Ben Abbas is acknowledged for his help with sequencing.

## Supplementary Materials

**Fig. S 1:**
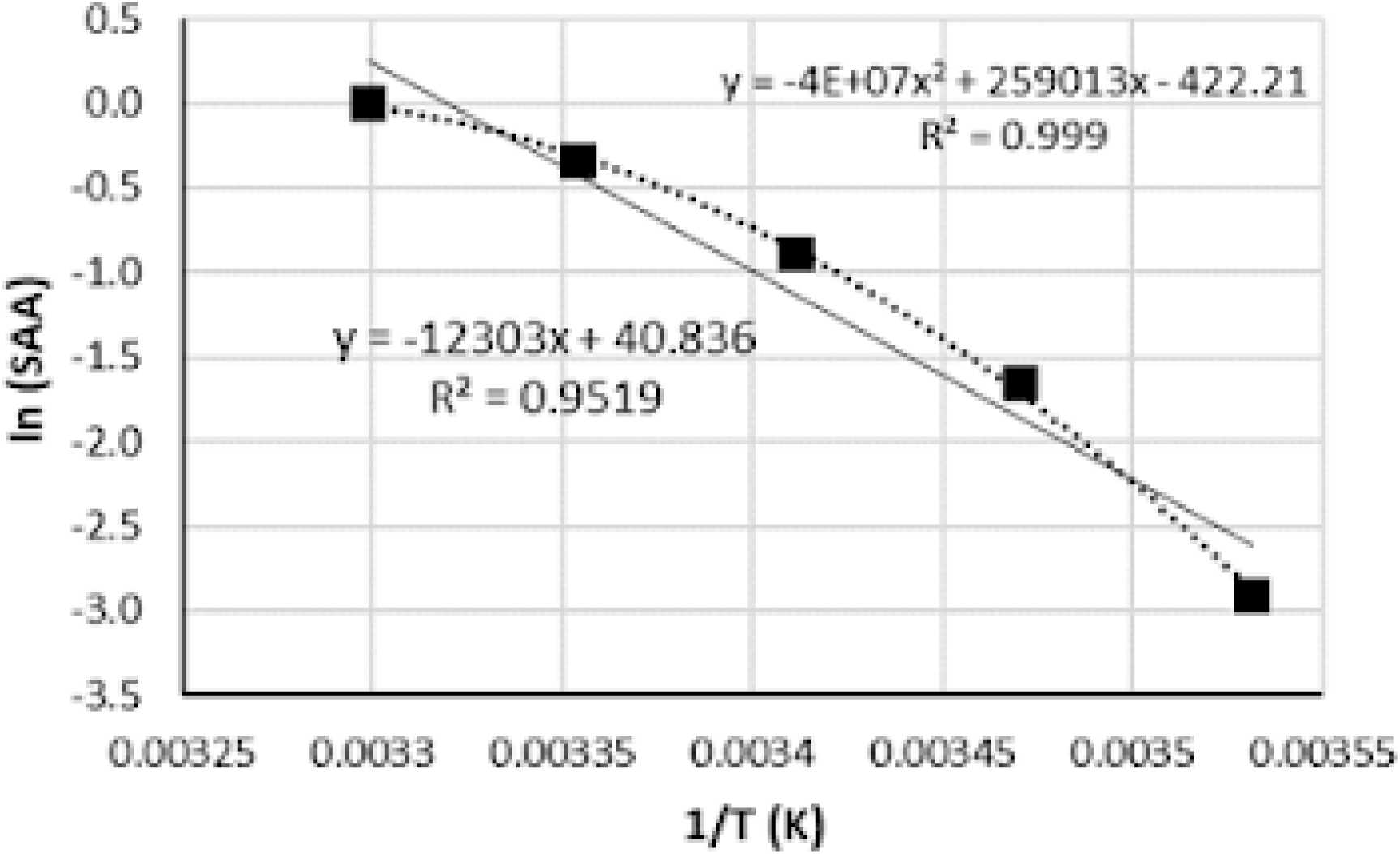
Arrhenius plot for anammox culture from Strass (Austria).

**Tab. S 1:**
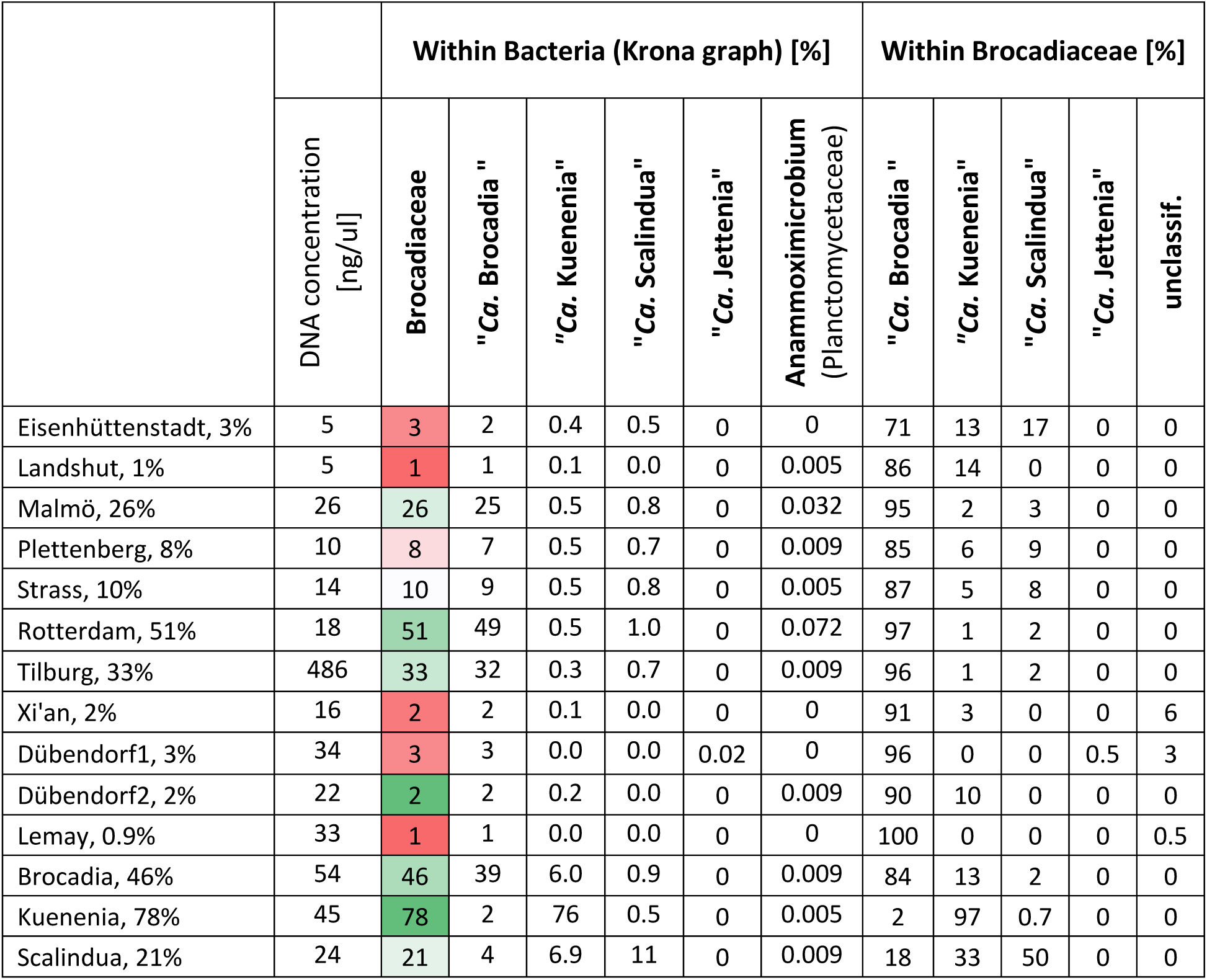
Results of 16S rRNA-gene amplicon sequencing in various anammox biomasses, including family Brocadiaceae, genera “*Ca*. Brocadia”, “*Ca*. Kuenenia”, “*Ca*. Scalindua”, and “*Ca*. Anammoximicrobium”. The abundance is expressed as % of OTU within Bacteria and within Brocadiaceae.

### Sequencing preparation

Total genome DNA from samples was extracted using CTAB/SDS method. DNA concentration and purity was monitored on 1% agarose gels. According to the concentration, DNA was diluted to 1ng/μL using sterile water. 16S rRNA/18SrRNA/ITS genes of distinct regions (16SV4/16SV3/16SV3-V4/16SV4-V5, 18S V4/18S V9, ITS1/ITS2, Arc V4) were amplified using specific primers (e.g. 16S V4: 515F-806R, 18S V4: 528F-706R, 18S V9: 1380F-1510R, et. al) with the barcode. All PCR reactions were carried out with Phusion® High-Fidelity PCR Master Mix (New England Biolabs). PCR products were mixed with the same volume of 1X loading buffer (contained SYB green) and loaded on 2% agarose gel for detection with electrophoresis. Samples with bright main strip between 400-450bp were chosen for further experiments. PCR products were purified with Qiagen Gel Extraction Kit (Qiagen, Germany). The libraries generated with NEBNext® UltraTM DNA Library Prep Kit for Illumina and quantified via Qubit and Q-PCR, were analysed by Illumina platform.

### Sequencing data processing

Paired-end reads was assigned to samples based on their unique barcode and truncated by cutting off the barcode and primer sequence. Paired-end reads were merged using FLASH (V1.2.7, http://ccb.jhu.edu/software/FLASH/, (Magoč and Salzberg 2011)). Quality filtering on the raw tags were performed under specific filtering conditions to obtain high-quality clean tags (Bokulich et al. 2013) according to the Qiime (V1.7.0, http://qiime.org/scripts/split_libraries_fastq.html, (Caporaso et al. 2010)) quality controlled process. The tags were compared with the reference database (Gold database,http://drive5.com/uchime/uchime_download.html) using UCHIME algorithm (UCHIME Algorithm,http://www.drive5.com/usearch/manual/uchime_algo.html (Edgar et al. 2011)) to detect chimera sequences. The chimera sequences were removed and the Effective Tags were obtained. Sequences analysis were performed by Uparse software (Uparse v7.0.1001 http://drive5.com/uparse/ (Edgar 2013)) using all the effective tags. Sequences with ≥97% similarity were assigned to the same OTUs. Representative sequence for each OTU was screened for further annotation. For each representative sequence, Mothur software was performed against the SSUrRNA database of SILVA Database (http://www.arb-silva.de/ (Quast et al. 2012)) for species annotations at each taxonomic rank (Threshold:0.8∼1). In order to study phylogenetic relationship of different OTUs, and the difference of the dominant species in different samples, multiple sequence alignments were conducted using the MUSCLE software (Version 3.8.31,http://www.drive5.com/muscle/ (Edgar 2004)).

**Fig. S 2:**
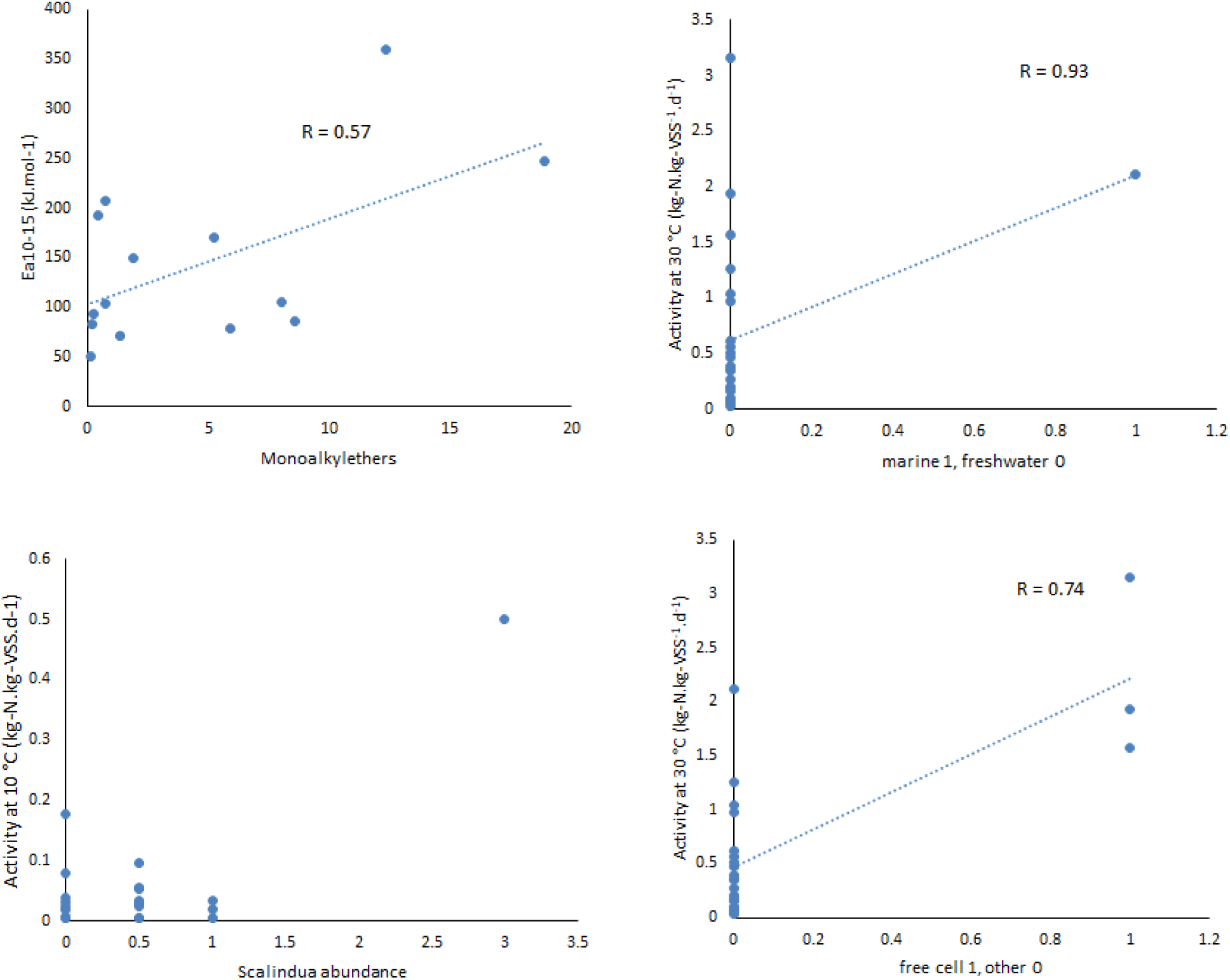
Correlations of selected process parameters and performance of anammox cultures, according to Pearson.

